# Effect of Disease Causing Missense Mutations on Intrinsically Disordered Regions in Proteins

**DOI:** 10.1101/2021.04.26.441554

**Authors:** Suryanarayana Seera, H.A. Nagarajaram

## Abstract

It is well known that disease-causing missense mutations (DCMMs) reduce the structural stability/integrity of the proteins with well-defined 3D structures thereby impacting their molecular functions. However, it is not known in what way DCMMs affect the intrinsically disordered proteins (IDPs) that do not adopt well defined stable 3D structures. In order to investigate how DCMMs may impact intrinsically disordered regions (IDRs) in proteins we undertook Molecular Dynamics (MD) based studies on three different examples of functionally important IDRs with known DCMMs. Our studies revealed that the functional impact of DCMMs is in reducing the conformational heterogeneity of IDRs which is intrinsic and quintessential for their multi-faceted cellular roles. These results are reinforced by energy landscapes of the wildtype and mutant IDRs where the former is characterized by many local minima separated by low barriers whereas the latter are characterized by one global minimum and several local minima separated by high energy barriers. Our MD based studies also indicate that DCMMs stabilize a very few structural possibilities of IDRs either by the newly formed interactions induced by the substituted side chains or by means of restricted or increased flexibilities of the backbone conformations at the mutation sites. Furthermore, the structural possibilities stabilized by DCMMs do not support the native functional roles of the IDRs thereby leading to disease conditions.

## Introduction

In the recent past, many proteins referred to as intrinsically disordered proteins (IDPs) have been discovered (Uversky & Dunker, 2010; Uversky, Gillespie, & Fink, 2000; Wright & Dyson, 1999). IDPs are those proteins which do not adopt stable 3D structures under physiological conditions yet are involved in certain critical cellular functions such as, cell signaling, cellular recognition (Uversky, Oldfield, & Dunker, 2005), regulation of transcription and translation (Dunker, Brown, Lawson, Iakoucheva, & Obradovic, 2002; Obradovic et al., 2008) etc. IDPs are also known to be cancer-associated proteins (Iakoucheva, Brown, Lawson, Obradovic, & Dunker, 2002). The intrinsically disordered regions (IDRs) in IDPs facilitate easy access to post-translational modifications such as phosphorylation and ubiquitination (Iakoucheva et al., 2004; Radivojac et al., 2011). Functional yet not adopting stable 3D structures, has challenged the traditional sequence-structure-function paradigm that necessitates stable 3D structure as a prerequisite for protein function. Although IDPs do not adopt stable 3D structures it is believed that they adopt multiple interconverting conformational states that help them to interact with multiple partners and hence display multiple functions (Wright and Dyson 2015). IDPs/IDRs are characterized by distinct sequence composition enriched with charged, polar residues and Glycyl and Prolyl residues and are generally depleted with large hydrophobic residues and hence are not capable of folding spontaneously, instead, they exhibit conformational heterogeneity (Campen et al., 2008; Vucetic, Brown, Dunker, & Obradovic, 2003). Unlike the regions with stable and well defined secondary structures (ordered regions) the IDRs are not well conserved (Brown, Johnson, Dunker, & Daughdrill, 2011). While the energy surfaces of ordered proteins are characterized by a single well-defined global energy minimum, IDPs display energy surfaces with multiple local minima separated by low energy barriers and appear rugged (Fisher & Stultz, 2011; Schaefer, Schlessinger, & Rost, 2010). IDPs generally appear as highly connected hubs in protein-protein interaction networks (Dosztanyi, Chen, & Dunker, 2006; Haynes et al., 2006) and the disordered regions in them usually harbor Short Linear Motifs (SLiMs) that are involved in transient, reversible weak protein-protein interactions (Davey et al., 2012). The inherent conformational heterogeneity and the SLiMs enable IDPs to bind to multiple partners (Oldfield et al., 2008; Patil & Nakamura, 2006).

Despite having conformational heterogeneity in isolation, an IDP adopts discrete conformation upon interaction with its partner while executing its function, and can also adopt different conformations with different partners (Qin et al., 2005). There are two proposed models that explain the binding mechanism of IDPs. One model suggests that IDP adopts one of the conformational states suitable for the interacting partner. In other words, the bound form of an IDP pre-exists in the conformational ensemble and is stabilized by its partner. For example, the interaction between the p27kip1 and cdk2 cyclin (Borriello, Cucciolla, Oliva, Zappia, & Della Ragione, 2007) as well as the interaction between p53 and MDM2 (Espinoza-Fonseca, 2009; Kar et al., 2002; Lowry, Stancik, Shrestha, & Daughdrill, 2008) are the two well-established studies of conformational selection. Molecular dynamics simulations have revealed that the bound form of p27kip1 populates appreciably in their conformational ensemble and is stabilized by its partner. The second model referred to as “Coupled folding and binding”, indicates that an IDP binds to the partner in the disordered state and forms a transient complex that is stabilized by non-specific hydrophobic contacts. This wobble intermediate evolves as the native complex with stable intermolecular interactions (Dogan, Mu, Engström, & Jemth, 2013; Sugase, Dyson, & Wright, 2007; Turjanski, Gutkind, Best, & Hummer, 2008). These favourable intermolecular interactions reduce the entropy and drive IDPs toward a low energy conformational state.

The intrinsic disorder is not an uncommon feature in higher eukaryotic genomes. The total constitution of disorderedness in the known genomes varies from 1-18% (in archaea and bacteria) to 25-41% (in eukaryotes) (Dunker, Obradovic, Romero, Garner, & Brown, 2000). It has also been reported that 40% of the human proteome is intrinsically disordered (J. J. Ward, Sodhi, McGuffin, Buxton, & Jones, 2004) and hence there is a possibility that some of the reported nonsynonymous disease causing missense mutations (DCMMs) map on to the IDRs of IDPs (Uversky, Oldfield, & Dunker, 2008; Vacic & Iakoucheva, 2012). However, it is not clear how a DCMM affects an IDP/IDR. It is known that DCMMs destabilize 3D structures of ordered proteins thereby rendering them non-functional. How does a DCMM then affect a protein which is already unstructured? In order to investigate into this question, we carried out molecular dynamics simulations on IDRs carrying DCMMs. Our studies revealed that disease causing missense mutations reduce the inherent conformational heterogeneity of IDRs that underpins their cellular functions.

## Materials and methods

In order to select IDRs harbouring DCMMs, we first prepared a list of human proteins harbouring IDRs of lengths ≥ 30 residues. The IDRs were predicted using DISPROT3 (Jones & Cozzetto, 2015). This was followed by a search for their known 3D structures (as complexes) in the PDB databank with an idea of obtaining 3D structures for using them as starting points for MD simulation studies. Known disease causing missense mutations were retrieved from humsavar (www.uniprot.org>docs>humsavar) and humvar (ftp://genetics.bwh.harvard.edu/pph2/training) datasets. These steps are illustrated in supplementary figure 1. For the MD studies we selected three functionally important IDRs harboring DCMMs. The IDRs are: a) B-chain of 1CEE (Dye-terminator et al., 1999) (henceforth referred to as CRIB IDR, encompassed between the 230 to 247 residues) b) A-chain of 1WLP (Ogura et al., 2006) (henceforth referred to as PRR IDR, encompassed between the 148 to 167 residues) and c) H-chain of 3TWR (Guettler et al., 2011) (henceforth referred to as TRM IDR, encompassed between the 410 to 425 residues). The TRM IDR (H-chain of 3TWR) had a missing region in its X-ray structure and this region was modelled using the Loop modelling tool available in MODELLER (https://salilab.org/modeller/9.25/) and procheck (https://oc.ebi.ac.uk/) used to validate the stereochemical properties.

The wild-type (WTs) IDRs extracted from the PDB files were subjected to energy minimization. The energy minimized structures were used to produce their disease mutant forms (MTs) by means of replacing the WT residues with the MT residues. By this, we obtained 5 MTs from 3 WTs and they are: one each for CRIB IDR and PRR IDR and three for TRM IDR. The residue replacements were carried out using pymol (http://pymol.org/educational/v1.74).

### MD simulations

MD simulations were carried out using GROMACS version 5.12 (http://www.gromacs.org/Downloads) with charmm36-nov2018.ff force field for the energy calculations. It may be noted that charmm36-nov2018.ff is the force field that has been developed for simulations of IDPs (Huang et al., 2017). SPCE water models were used for solvation with 1.0 nm periodic boundary conditions. Steepest descent minimization algorithm was used to perform energy minimization to remove any steric clashes and bad geometries associated with the selected structures. The termination criteria for minimization was ≤ 1000 KJ/mol. The system was equilibrated in terms of temperature and pressure at their reference values by performing the NVT followed by NPT ensembles. During the equilibration, atomic positions in IDRs were restrained. Temperature and pressure maintained constantly by applying the modified Berendsen thermostat algorithm and Parrinello-Rahman barostat algorithm respectively. LINCS algorithm was used to constrain the covalent bonds. A distance cut-off of 1nm was used to detect short-range electrostatic interactions and van der Waal interactions and were modelled by a varlet algorithm cut-off-scheme. Particle Mesh Ewald was used to calculate the long-range electrostatic interactions. After equilibration, MD simulations were performed for 100ns by constraining the terminal residues. MD simulations were carried out on a Super Microcomputer with GPU Tesla card K40.

### Analysis of MD simulations

We used GROMACS gmx-toolbox to analyze properties such as RMSD, RMSF, the radius of gyration, and solvent accessibility surface area, clustering, analysis of intramolecular hydrogen interactions and secondary structural analysis etc. Free energy landscapes of the IDRs were computed by dihedral principal component analysis (Michel Espinoza-Fonseca, Ilizaliturri-Flores, & Correa-Basurto, 2012) and the FEL plots were plotted by using the mathmatica software (https://www.wolfram.com/mathematica/).

## Results and discussion

### Selection of IDRs for MD simulations

About 1766 DCMMs mapped on to the 771 IDRs of 565 proteins. These proteins belong to different functional classes that include signaling proteins, proteins involved in transcription and translation and other functional groups (Supplementary figure: 2) indicating that the DCMMs have been distributed in IDRs of many functional classes of proteins. Our search for IDRs with known PDB structures (as part of complexes) harboring mutational sites of known DCMMs yielded 111 IDRs with 275 DCMM mutational sites. Of these IDRs we selected only those with minimum length of 15 residues harboring at least one experimentally validated Eukaryotic Linear Motif (ELM) involved in PPI as evident from the availability of complex structure in PDB. This ensured that the IDR that we are selecting is functionally important for the protein and hence any mutation that affects its structure would potentially affect its functional role and hence render the protein with an impaired function. We selected three IDRs for detailed MD simulations. The details of these IDRs are given below.

a. The CRIB IDR) is located between the residues 230 to 247 in the GTPase binding domain (GBD) of the Wiskott Aldrich Syndrome Protein (WASP) and forms a complex with the Cdc42 in order to stimulate the actin nucleation and polymerization. This 18 residue long IDR with the known NMR structure (B chain in 1CEE; Dye-terminator et al., 1999) is part of a long IDR (148 to 247). The reported DCMM Ala236Glu is located near to the N terminal side of the CRIB motif (238 to 247) in GBD of WASP. This mutation is likely to induce new Glu side chain H-bonding interactions with its nearby amino acids.
b. The PRR IDR located between the residues 148 to 167 in the cytosolic domain of membrane bound p22^phox^, is a part of long IDR located between the amino acid residues 138 and 180. The proline rich region (PRR) from 151 to 160 residues is very crucial for the interaction with the SH3 domain of cytosolic p47^phox^ in phagocyte NADPH oxidase complex (Ogura et al., 2006). Upon this interaction, the NADPH oxidase is activated and generates the reactive oxygen species (ROS) to kill the infected microbes. The DCMM Pro156Gln is situated within the PRR. The Pro residue when substituted by Gln certainly increases the backbone flexibility at the mutation site and also likely to induce new polar interactions. An NMR structure (148 to 167 residues of A chain of 1WLP; Ogura et al., 2006) is available for the PRR region which was used for the present study.
c. The TRM IDR is the tankyrase binding motif encompassed between the 410 to 425 residues and is a part of the long IDR (157 to 445) of the SH3 binding protein2. A crystal structure (H-chain of 3TWR; Guettler et al., 2011) is available for the segment between 410 and 425. It has been shown that, the hexapeptide (415 to 420) present in the TRM IDR acts as the recognition site for the tankyrase2, a member of poly(ADP-ribose) polymerase (PARP) family protein in order to regulate 3BP2 in terms of stability and degradation (Guettler et al., 2011). The DCMMs Arg415Pro, Pro418Leu and Gly420Glu are located in the hexapeptide. Arg to Pro at 415 position replaces the charged residue Arg to Pro that induces conformational restriction at the mutation site whereas the DCMM Pro418Leu increases the conformational flexibility. The DCMM Gly420Glu decreases the conformational flexibility and also likely to induce new interactions potentially made by the polar Glu with the nearby amino acid residues.

As can be seen the DCMMs alter conformational flexibilities and are likely to induce new interactions.

### MD Simulations

As already mentioned in the Methods section, the MT structures were generated by replacing the wild-type residues with the mutant residues and, where required, the side chain conformations were set to their best rotameric states with no steric clashes with the rest of the structure. Both the WT and MTs were subjected to the MD simulations for 100 ns as detailed in the methods section. The information of the molecular system which includes the number of residues, number of atoms, number of solvent molecules and number of energy minimization steps are summarized in Supplementary table 1.

The RMSD trajectories reveal how structure changes with respect to the starting structure under the influence of interatomic interactions during the entire course of MD simulation. The RMSD plots as well as the box plots shown in Figure:1 reveal that, in general, the MTs undergo significantly larger conformational changes from their starting structures as compared with their WTs during the simulations. It is clear that the RMSD values corresponding to the MTs are significantly higher than the WTs.

**Figure 1:**
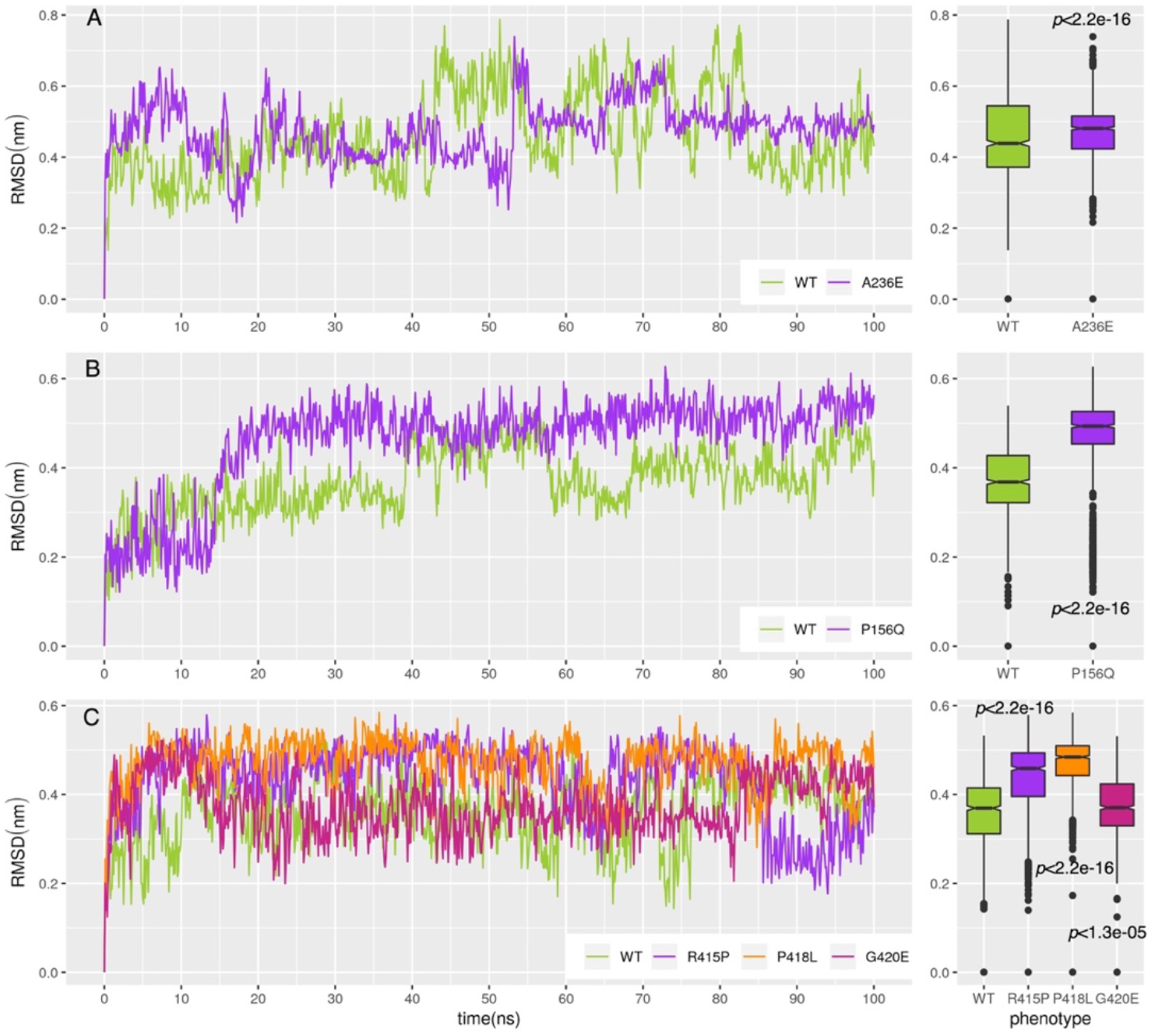
RMSD trajectories for the wild-type (WT) and mutant (MT) structures of (A) CRIB IDR, (B) PRR IDR and (C) TRM IDR. The adjacent panels give the box plots of the RMSD values for the WT and MT structures during the entire MD simulation period after the equilibration. It can be seen, in general, the RMSD values are significantly higher in MTs as compared with WTs indicating that the structural deviation of MTs, from the starting structure, are higher as compared with WT. The *P*-values were calculated by the Kolmogorov–Smirnov test.

The radius of gyration (Rg) and solvent accessibility surface area (SASA) indicate compactness of the protein and contribution of the surface area accessible to the surrounding solvent respectively. Figures: 2 and 3 show the trajectories of these two parameters. In general, the MTs show lower Rg values as compared with their respective WTs (with the exception of one mutation Arg415Pro in TRM IDR) indicating the MTs adopt lesser compact structures and hence show lower solvent accessibilities than their respective WT counterparts.

**Figure 2:**
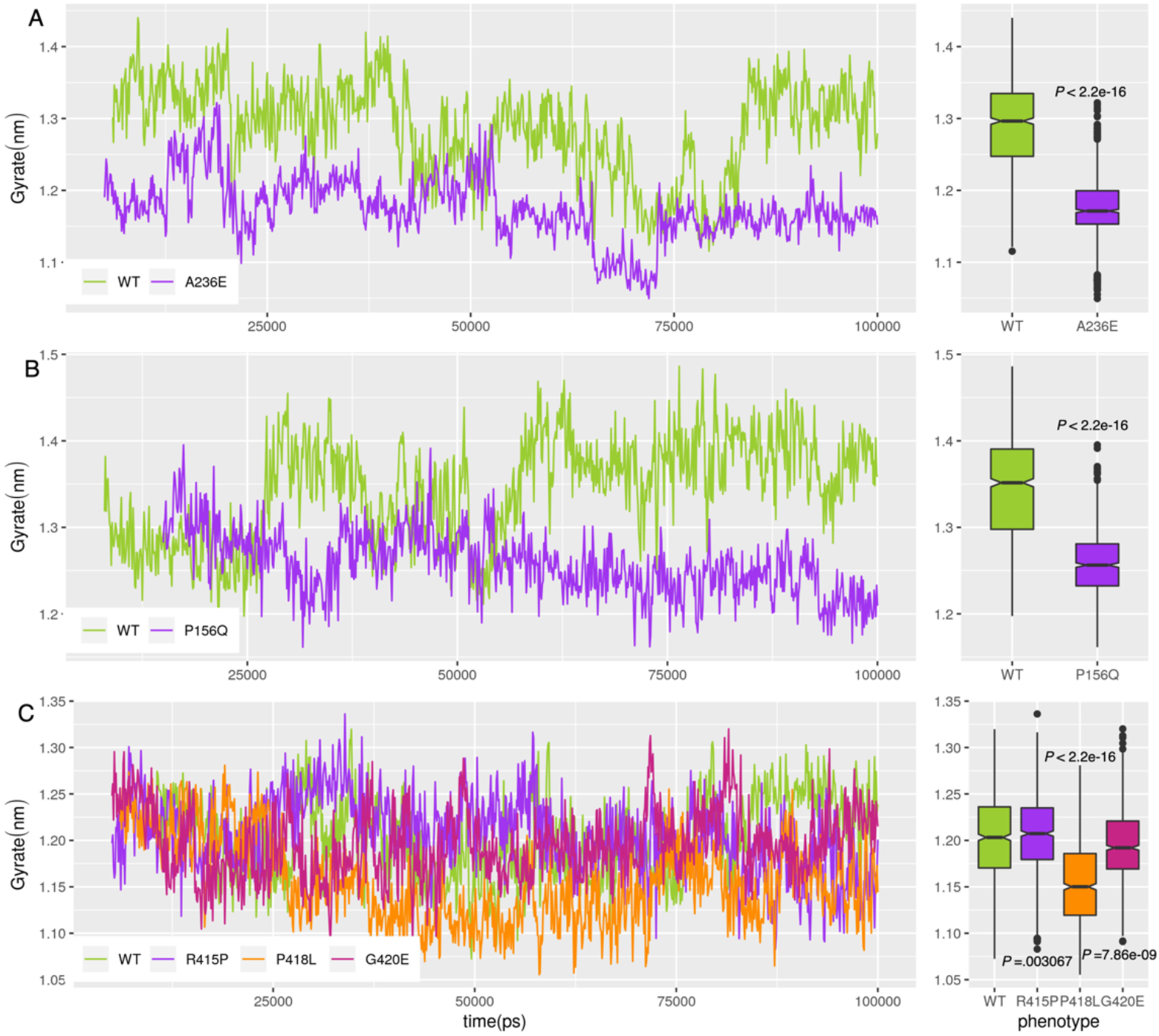
The trajectories of radius of gyration of (A) CRIB IDR, (B) PRR IDR and (C) TRM IDR. The adjacent box plots give the distribution of radius of gyration values in WT and MTs. It can be seen that the MT IDRs, in general, show lower values of radius of gyration as compared with their corresponding WTs, indicating that they adopt more closed conformations than their WTs. The *P*-values were calculated by the Kolmogorov–Smirnov test.

**Figure 3:**
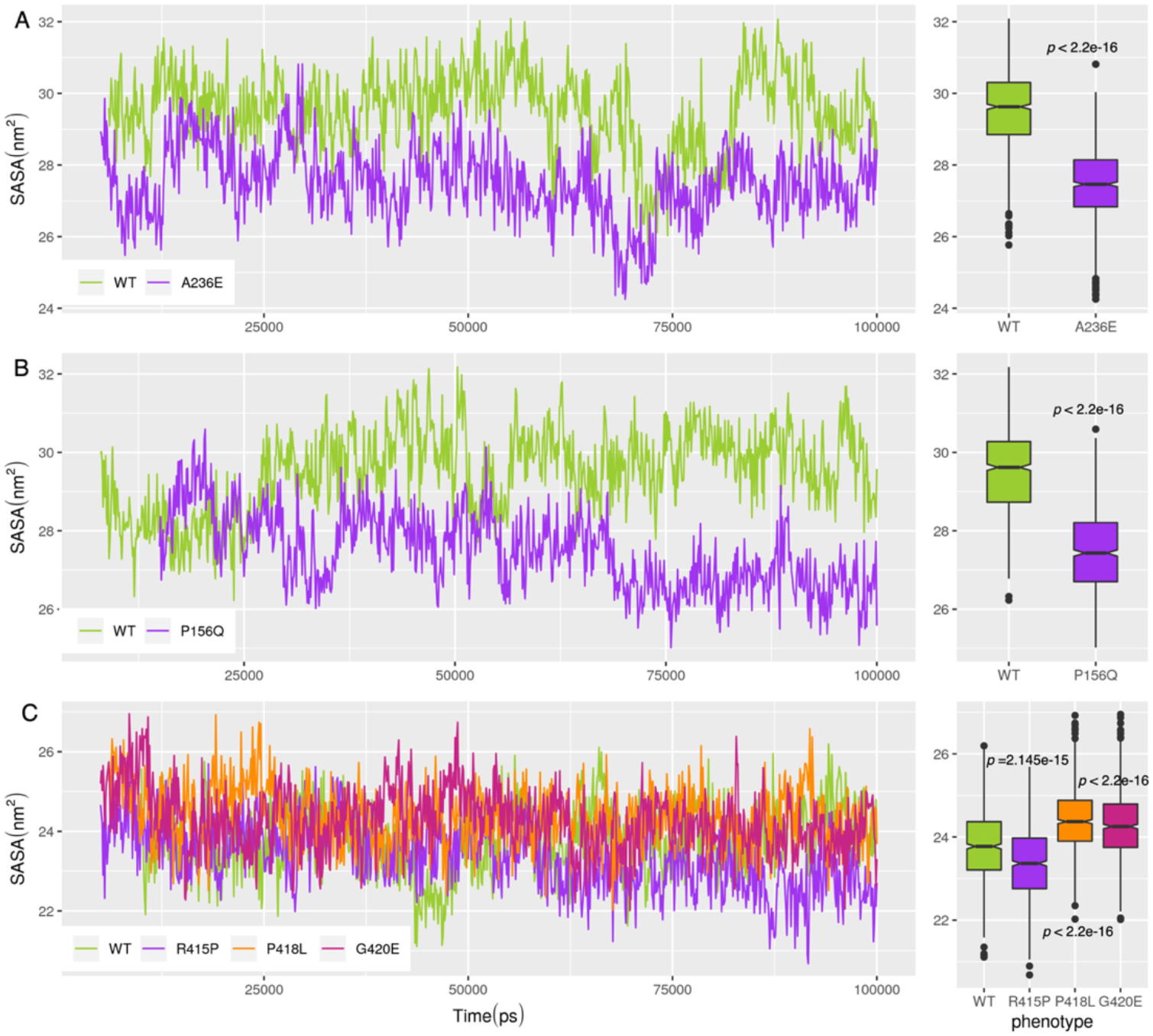
Trajectories of solvent accessibility surface area (SASA) of (A) CRIB IDR (B) PRR IDR and (C) TRM IDR and their distribution are shown in the adjacent box plots. SASA values are correlated with the radius of gyration values. In general, the MTs show significantly lower SASA values then their WTs indicating that they adopt structures that lower solvent accessible surfaces as compared with WTs. The *P*-values were calculated by the Kolmogorov–Smirnov test.

RMSF plots give insights into the flexibility of the regions with respect to the average structure of the trajectory during the simulations. The RMSF plots (Figure: 4) show that MTs display lower conformational flexibilities than their respective WTs except in the case of Pro418Leu in the TRM IDR.

**Figure 4:**
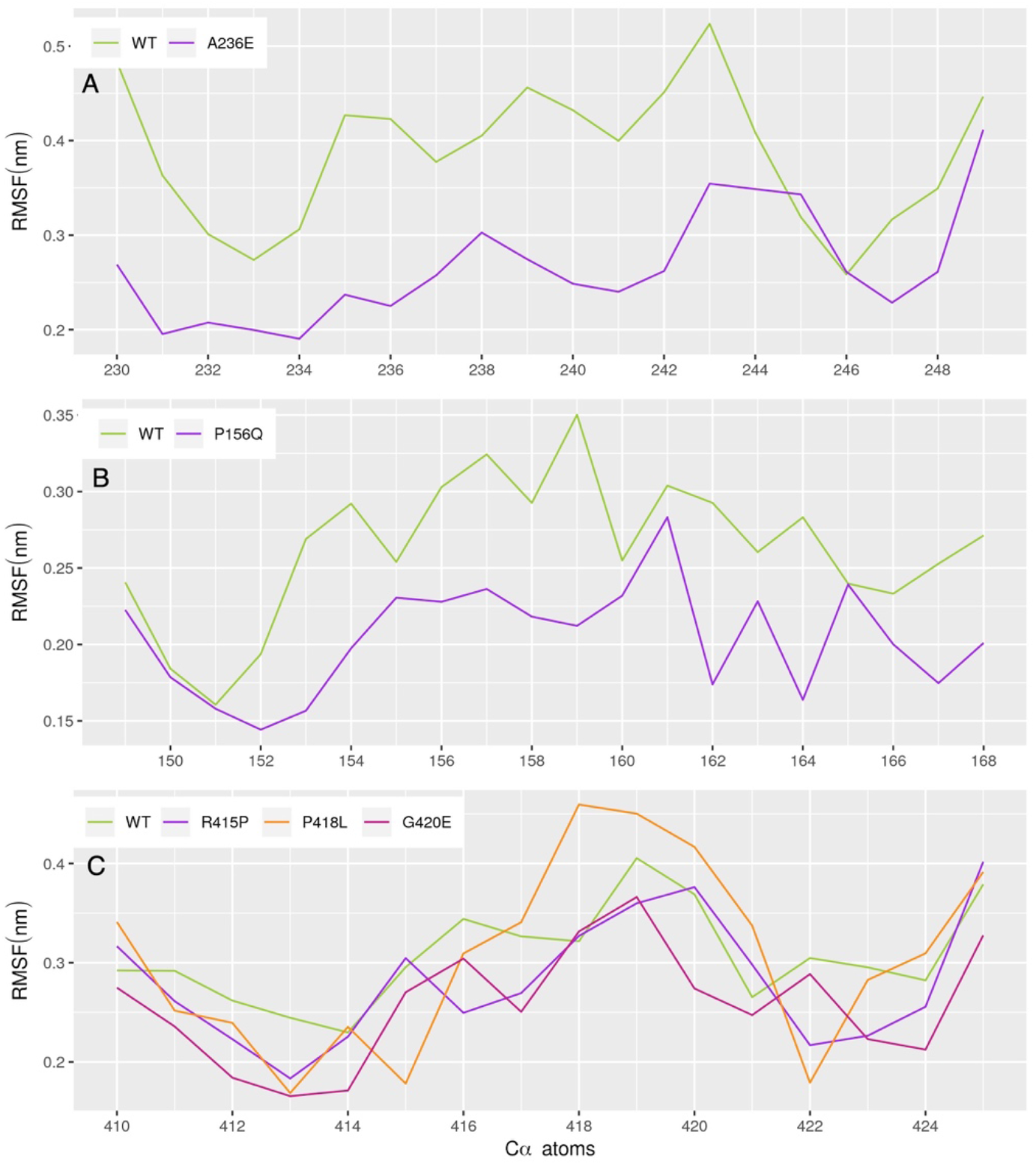
The RMSF values of (A) CRIB IDR (B) PRR IDR and (C) TRM IDR. Positional atomic fluctuations (local flexibility) are indicated by RMSF values computed at every amino acid residue. While computing RMSF values of the residues only Cα atoms were considered.

The aforementioned results indicate that, in general, IDRs harboring DCMMs adopt compact and less conformationally flexible structures that are highly deviated from their WTs. In order words, DCMSs seem to reduce the conformational heterogeneity of IDRs and make them to adopt a few conformational possibilities. This aspect was further investigated as given under.

### Analysis of distinct conformational states of IDRs and their disease mutant forms

To find the distinct conformations adopted by the WTs and their MTs, we subjected all the instantaneous structures captured during MD simulations to a clustering algorithm as discussed in Methods section. We considered the top 20 percentile of highly populated clusters as they constitute more than 60% of the snapshots and more than 70% of the simulation time. Each cluster can be considered as the representative of a distinct conformational state adopted by WT IDRs and their MTs during simulations and henceforth each cluster of snapshots is considered as a representation of a distinct conformational state adopted by IDR. Needless to mention, the number of distinct clusters found during a simulation can be related to the extent of conformational heterogeneity of the IDRs.

Our analysis revealed that the WT of CRIB IDR adopts 42 conformational states (i.e., it forms 42 clusters) comprising 62% of the snapshots that correspond to 58.4ns of simulation time. As compared with the WT, the corresponding MT forms only 10 clusters comprising of 70% of the snapshots that corresponds to 77ns of simulation time. In the case of PRR, the WT forms 13 clusters constituted by 80% of snapshots and corresponds to 73.9ns and the corresponding MT forms only 5 clusters constituted by 89.2% snapshots and of 75.9ns simulation time. In the case of TRM IDR, the WT forms 17 clusters constituted by 78.8% of snapshots and of 63.8ns of simulation time. Whereas the MT Arg415Pro forms 10 clusters constituted by 80.35% snapshots of 76.4ns, MT Pro418Leu forms 9 clusters made of 78.6% of snapshots of 77.3ns and the MT Gly420Glu forms 10 clusters constituted by 82.16% of frames of 74.8ns.

The above results indicate that, in general, WTs adopt several distinct conformations indicating their conformational heterogeneity whereas their MT counterparts do not adopt those many conformational states and seem to have ‘locked up’ in a few conformations (Figure: 5).

**Figure 5:**
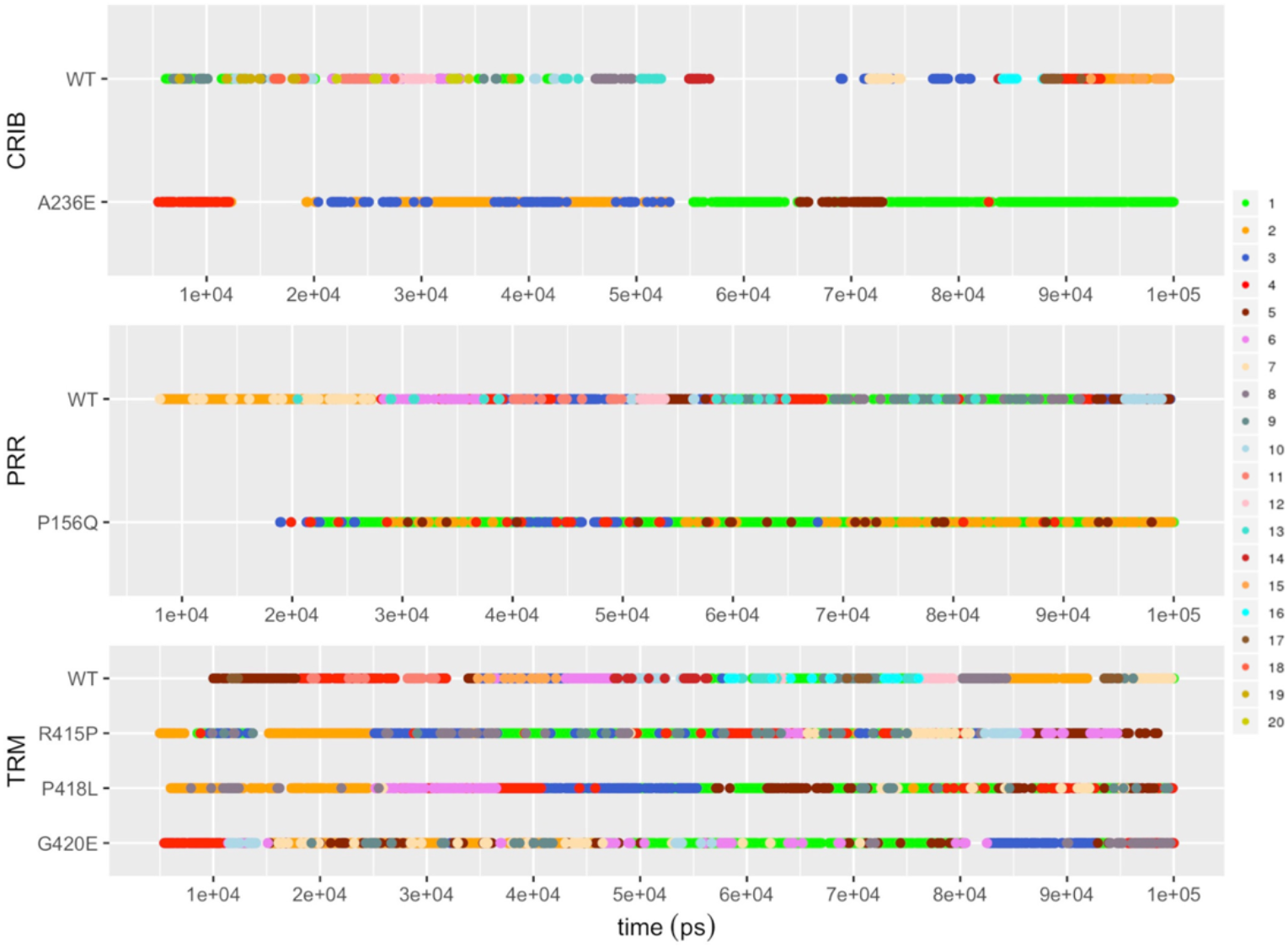
The resident times of unique conformational states of WTs and MTs. Each colour represents one unique cluster in a simulation and it should be noted that within an IDR or across IDRs having the same color does not indicate the equivalence in the concerned conformational state. The continuity of a colour represents the persistence of that particular conformational state over a period of time. Furthermore, more the number of times the same color dot appears but may be at different times, indicate the same conformational state is revisited those many times. For the sake convenience only the top 20 clusters are depicted in this diagram.

We examined the intramolecular hydrogen bonds in WT and MTs during simulations. The MTs are stabilized by a greater number of intramolecular hydrogen bonds as compared with WTs (Figure: 6). This indicates that the mutations induce new H-bonding interactions that stabilize their conformations. This is the reason why very few conformational states are adopted by MTs during the simulations and hence explains the lower conformational heterogeneity shown by MTs as compared with their WTs.

**Figure 6:**
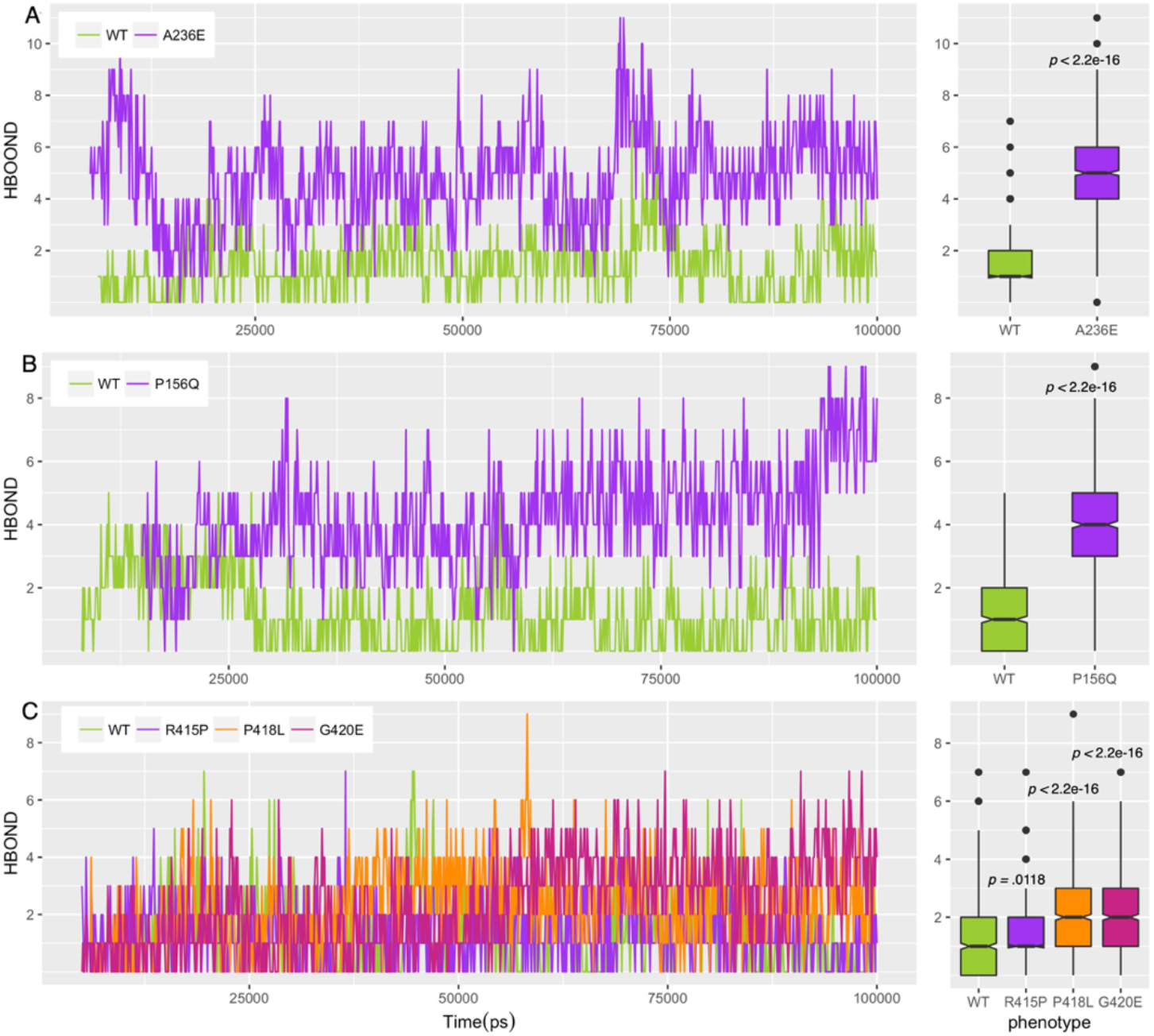
The number of intramolecular hydrogen bonds detected in WTs and MTs in (A) CRIB IDR (B) PRR IDR and (C) TRM IDR. Adjacent box plots give the distribution of the numbers of hydrogen bonds. In general, MTs form a significantly higher number of intramolecular hydrogen bonds than their WTs indicating higher stability of MT structures than their WT counterparts. The *P*-values were calculated by the Kolmogorov–Smirnov test.

**S**econdary structural analysis revealed that MTs adopt more often the short regular secondary structures such as a short helix and/or a beta strand as compared with the corresponding WTs which mostly adopt transient coils, turns and bends (Figure: 7a, 7b, and 7c). Needless to mention, the short secondary structures are stabilized by the hydrogen bonds formed in MTs that are not seen in the corresponding WTs (please see supplementary figures: 3a, 3b, 4a, 4b, 5a, 5b, 5c, and 5d).

**Figure 7a:**
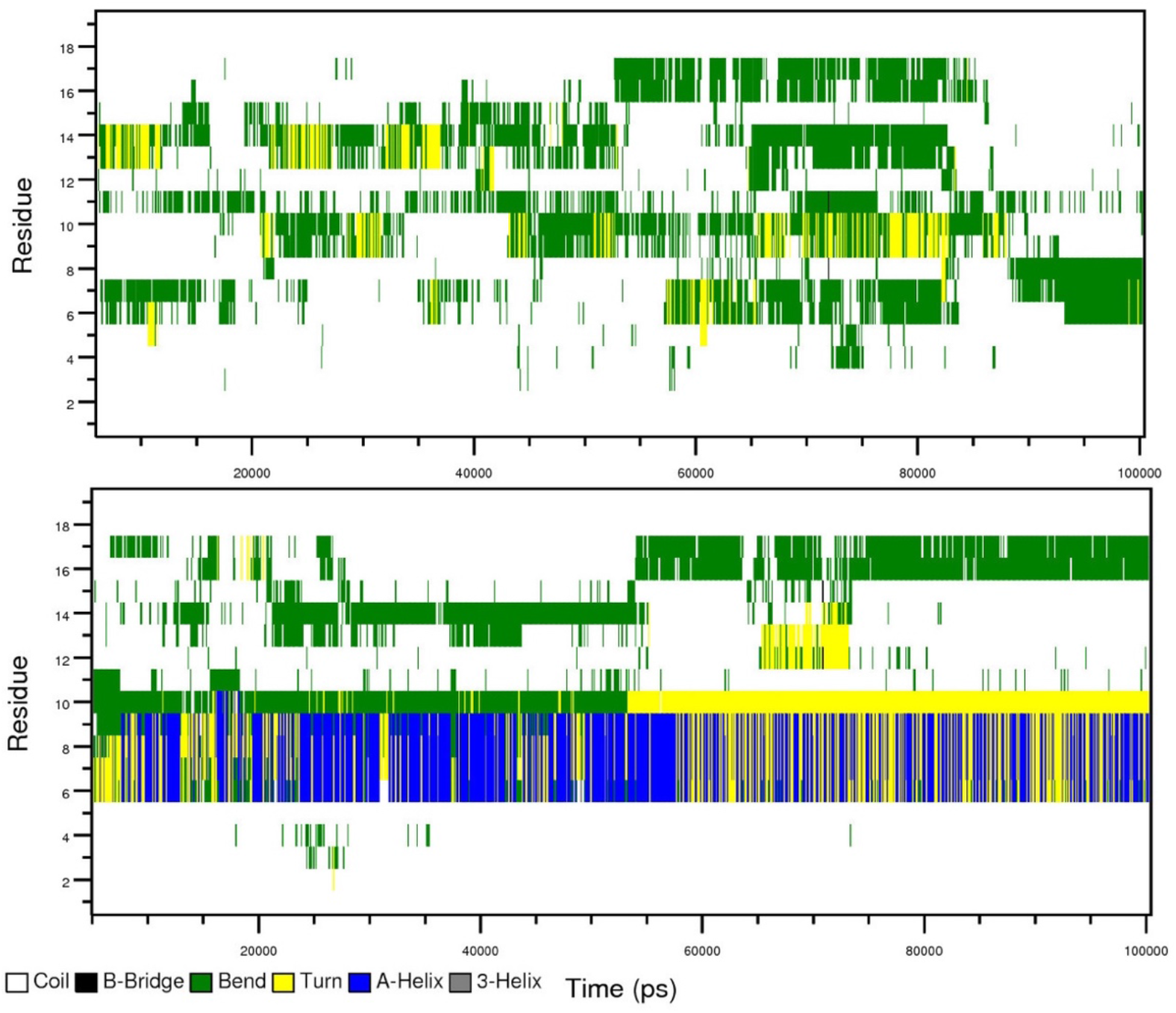
The WT IDR (top) of the CRIB motif visited several transient turns, bends and coils during the simulations. Whereas the MT IDR (bottom) visited stable α-helix throughout the simulations.

**Figure 7c:**
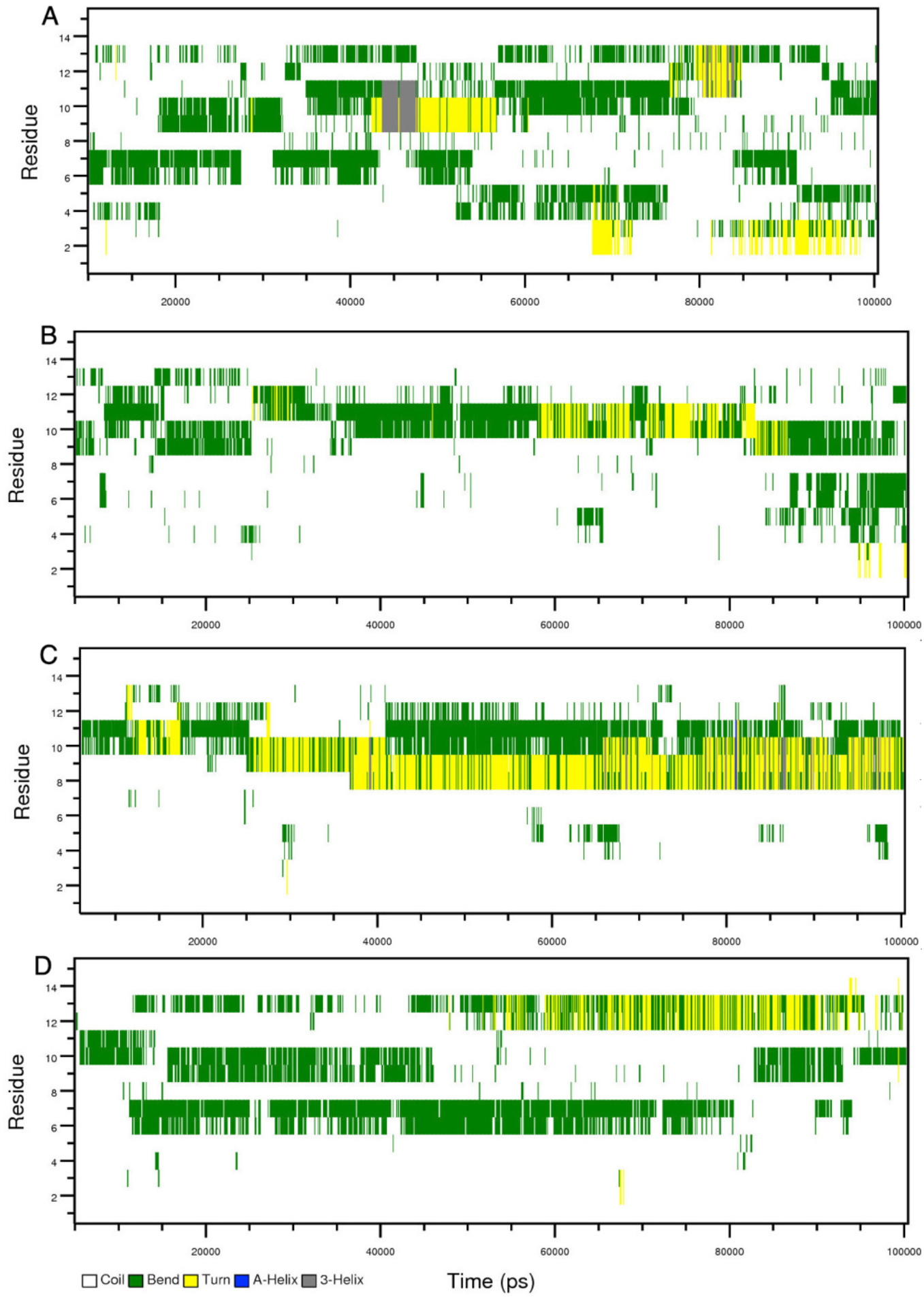
Bin A represents the SSEs of WT TRM and the rest of the bins B, C, and D represent the SSEs of MT IDRs Arg415Pro, Pro418Leu and Gly420Glu respectively. The WT mostly adopts transient coils, turns, 3-helix, and bends while the MT adopts continuous turns and bends during the MD simulations.

### Free energy landscapes of WT and MT IDRs

We constructed the free energy landscapes (FEL) for WTs and MTs by projecting the first two principal components (PCs) derived from the dihedral Principal Component Analysis (dPCA) (Figure 8a, 8b). In general, all the MTs exhibit fewer local minima as compared with their respective WTs correlating with the results obtained from the cluster analysis. The other important aspect is the widths and depths of the energy minima in MTs and WTs. In general, the MTs show wider and deeper energy wells than their WTs. The top 20 percentile distinct conformational substates from all the simulations of IDRs were mapped onto their respective FELs and found that the centroids of WT IDRs are located at different local minima (Supplementary figure 6, 7 and 8). In contrast to the WT IDRs, the centroids of respective MT IDRs are located at one or two local minima with the exception of MT IDRs such as Arg415Pro and Pro418Leu of TRM IDR.

**Figure 8a:**
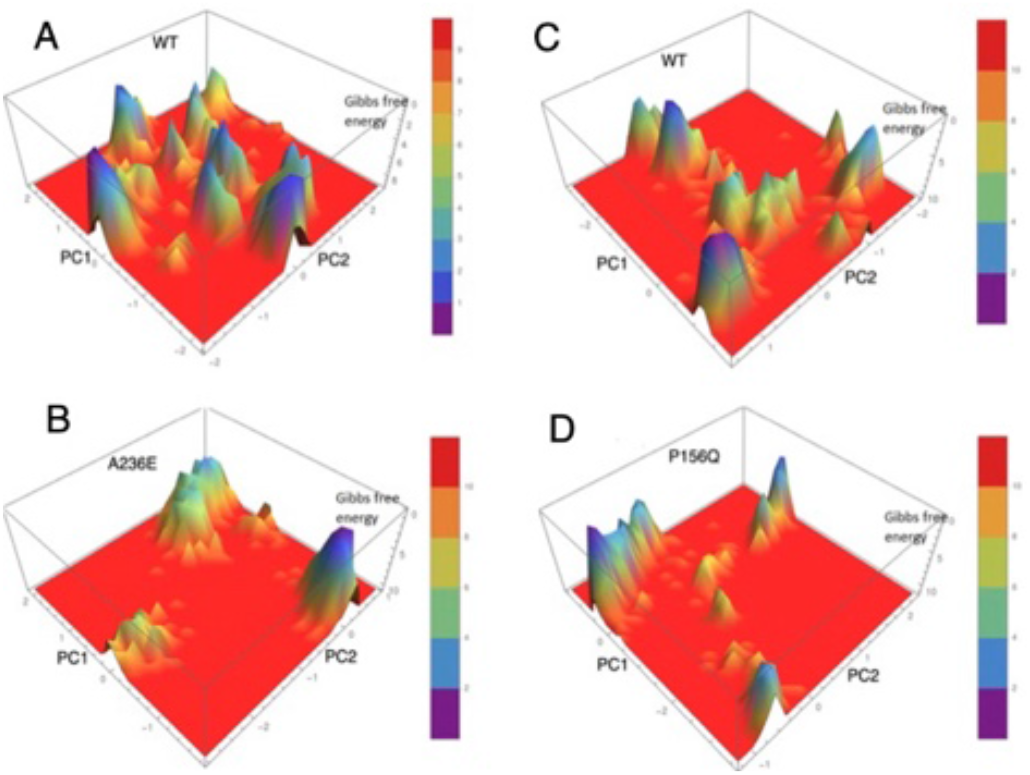
Free energy landscapes of IDRs of CRIB and PRR motifs. A and B represent the WT and MT of CRIB IDR respectively. C and D represent the WT and MT of PRR IDR respectively. PC1 and PC2 represent the principal components and which were obtained from the dPCA method (please see methods section). The scale represents the Gibbs free energy. The FEL of WT IDRs from both motifs shows more numbers of funneled like structures (local minima) than their respective MT IDRs. Due to the high number of local minima, the conformational space of WT IDRs appeared like uneven hilly FELs with several free energy basins, which is the characteristic of IDPs. The MT IDRs failed to exhibit several numbers of local minima in their FELs.

**Figure 8b:**
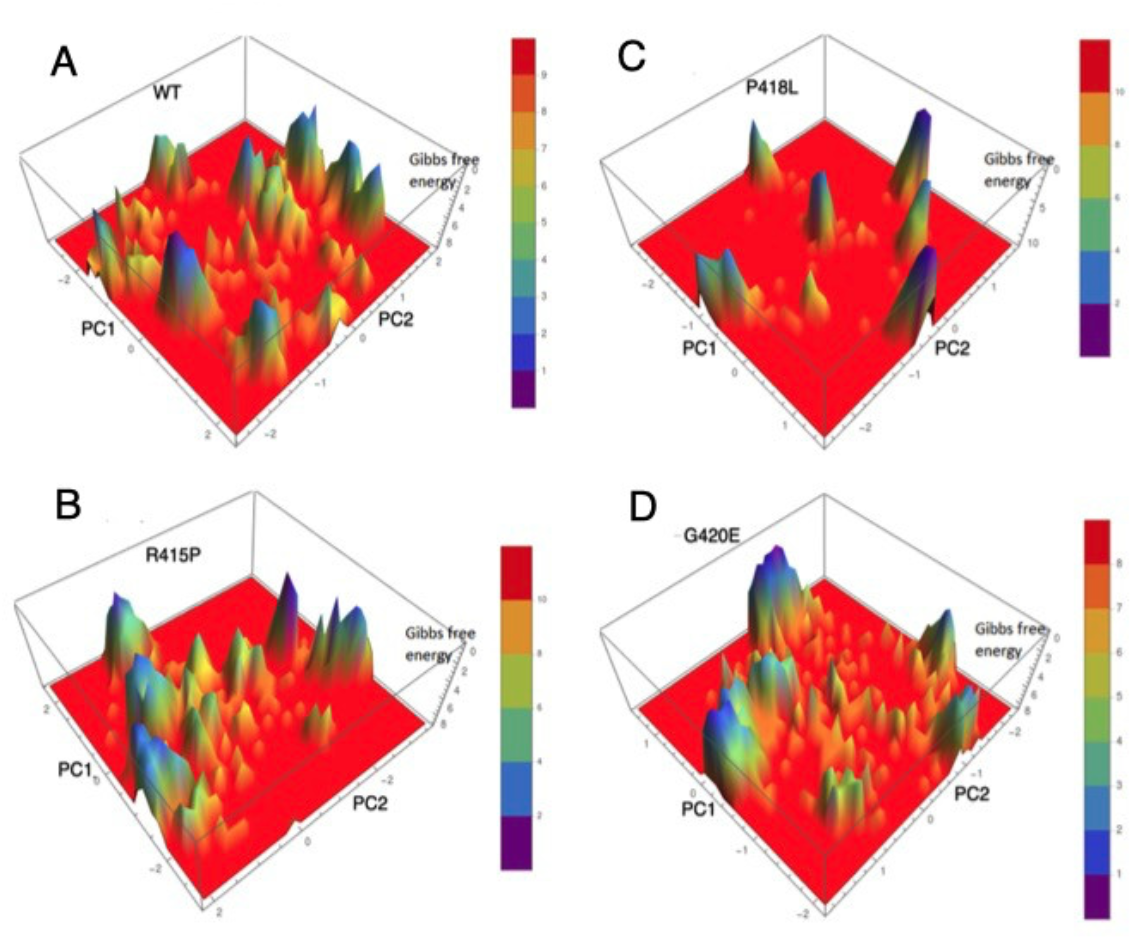
Free energy landscapes of IDR of TRM. The A bin represents the WT IDR and the rest of the three bins B, C, and D represent the MT IDRs R415P, P418L and G420E respectively. The WT IDR showed the multiple energy basins whereas the R415P and G420E IDRs showed some extent of ruggedness in their FELs, but the higher broadness of the local minima of MT IDRs indicated that most of the conformational substate during the simulations restricted to the one or two regions.

### The Disease Causing Mutations and their functional impact

In this section we discuss the IDRs and possible impact of the DCMMs on their structures and hence function.

The CRIB IDR is located between the residue numbers Lys230 to Val247 in the GTPase binding domain (GBD) of Wiskott aldrich syndrome protein (WASP). When Cdc42 binds to the CRIB IDR of GBD, the GBD displaces from the VCA (Verprolin homology, Cofilin homology, Acidic) region of the same protein in order to stimulate the actin nucleation and polymerisation along with the arp2/3 actin-nucleating complex (Cory, Cramer, Blanchoin, & Ridley, 2003).

As already mentioned, WT CRIB adopts 42 distinct conformations. In contrast, the MT CRIB adopts 10 distinct conformations. Our secondary structural analysis revealed that the N terminal region of the MT CRIB adopts a short helical structure and the mutated residue Ala236Glu stabilizes the helix with its new side chain intramolecular interactions (Supplementary figure: 3b) whereas WT adopts transient turns and bends. It can be inferred that the alpha helix formation stabilized by the mutation Glu at 236 position interferes with the interaction between the CRIB of GBD and Cdc42. As a consequence, the displacement of the GBD of WASP does not occur from the VCA region of the same protein and therefore the VCA region is not available to the Arp2/3 nucleation complex for binding. Absence of the interaction between the VCA of WASP and Arp2/3, the actin nucleation and polymerization does not occur in hematopoietic cells. As a result, the size of the platelets particularly decrease in the bloodstream and spleen destroys these platelets. Therefore, the reduction in number of the platelet cells in the bloodstream takes place and this condition is termed as thrombocytopenia 1 (Luthi, Gandhi, & Drachman, 2003).

The PRR IDR is located between the 148 to 167 residues in the cytosolic domain of the membrane bound p22^phox^ of the phagocyte NADPH complex. The proline rich region (151 to 160) of the PRR IDR adopts the polyproline type II helix (PPII) which is pivotal for the interaction (Ogura et al., 2006) and the IDR interacts simultaneously with the two tandem SH3 domains of cytosolic p47^phox^ of the same NADPH oxidase complex. This interaction activates the NADPH oxidase to generate the reactive oxygen species (ROS) in order to kill the infected microbes (Ogura et al., 2006). The PRR is conformationally constrained due to the presence of prolines in the sequence and the dihedral angles of the residues from the WT and MT of PRR IDR show similar distribution during simulations (Figure 9). The mutation occurs between two proline residues and hence the mutated residue Pro156Gln is also restricted to the poly proline type II helix conformation. However, it is interesting to find that Ser153 in MT adopts one distinct conformation corresponding to the β region in the Ramachandran Map whereas in WT (Figure 9)- it adopts two conformations-one in the β region and the other in right-handed α-region. This is perhaps due to the fact that in MT there is an increase in the hydrophilicity of the peptide as a consequence of missense mutation Pro156Gln which can change the water structure in the vicinity of the mutation site which in turn can create a water mediated interaction with the -OH group of Ser at 153 stabilizing that residue to just one conformational region in the Ramachandran Map. Therefore, it can be concluded that the DCMM Pro156Gln disrupts the PPII which is optimal for the binding with the SH3 domain of the partner protein. The absence of PPII leads to the lack of interaction between the cytosolic PRR domain of membrane p22^phox^ and the SH3 domain of cytosolic p47^phox^. The NADPH oxidase with no interaction between the p22^phox^ and p47^phox^ unable to generate the ROS upon microbial infection. This condition leads to a disease known as a chronic granulomatous disease (CGD) (Dinauer et al., 1991).

**Figure 9:**
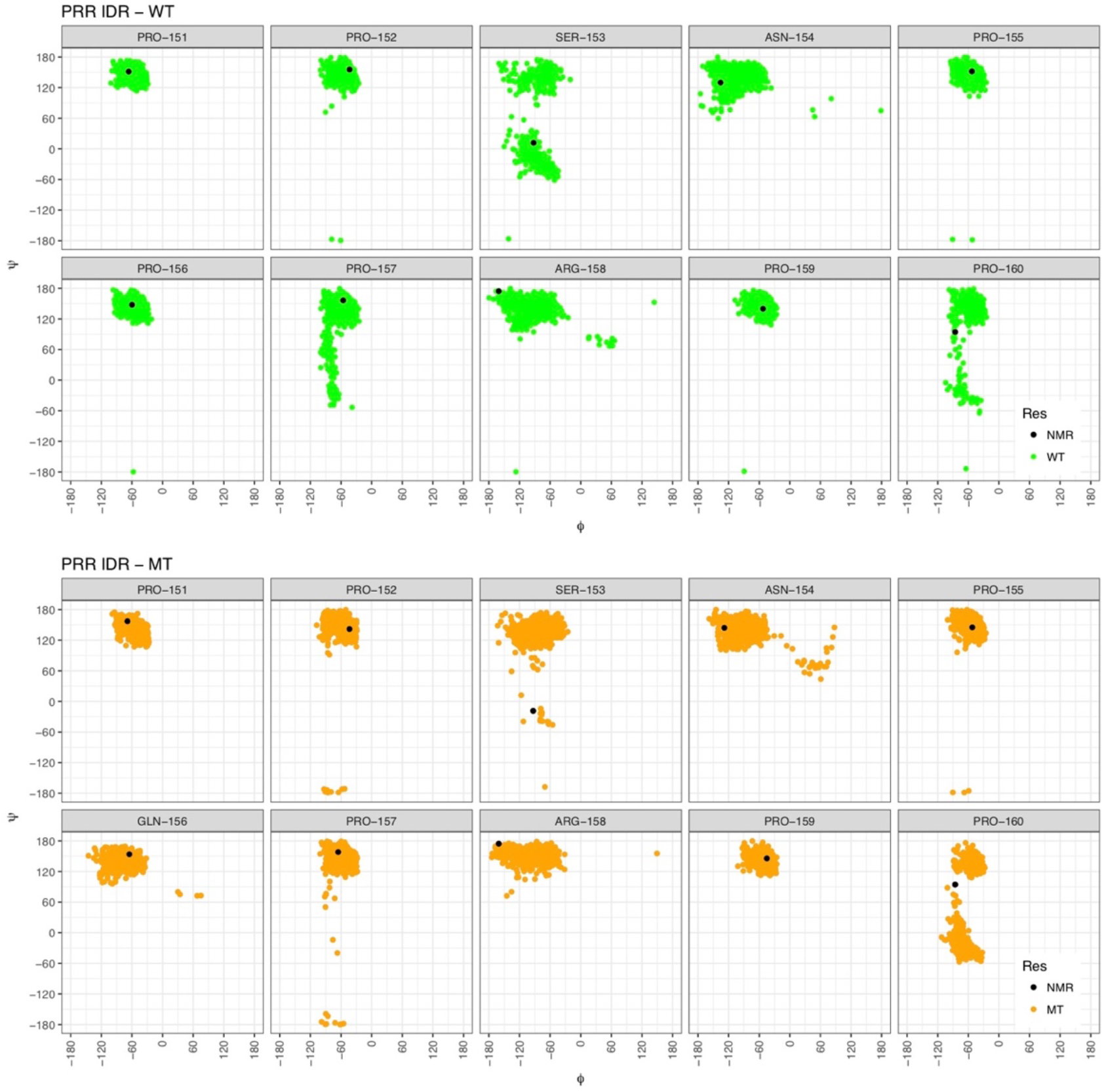
The dihedral angles of the proline rich region (151 to 160; involves in the binding) of the PRR IDR. The WT(PRO-156) and MT(GLN-156) residues showed the same distribution of the phi psi angles. But the residue SER-153 showed different distributions in MT as compared to the WT IDR during the simulations. SER-153 in the MT IDR almost not distributed in the right handed alpha helix region but the NMR SER-153(black) is located in this region. The residues other than SER-153 display same distribution in both the WT and MT IDRs.

The crystal structure depicts the TRM of 3BP2 binding to the tankyrase2 in an extended conformation (Guettler et al., 2011). It is clearly observed that conformation is well conserved during the MD simulations of WT IDR whereas such a structure is not adopted by the MT IDRs (Please see the RMSD values between the crystal structure and the distinct conformations of the WT and MT in Supplementary table-2). In other words, the mutations Arg415Pro and Pro418Leu perturb the extended conformation of the hexa peptide of 3BP2 and do not allow it to bind with the polyribosylation enzyme, tankyrase2. In the case of Gly420Glu of TRM IDR, the hexa peptide of four distinct conformational substates (IDR1, IDR4, IDR6, IDR8) have the RMSD values less than 0.1nm with respect to the hexa peptide of IDR of crystal structure (Supplementary table-2). Except IDR4, the mutated reside Glu420 in other three distinct conformations (IDR1, IDR6, and IDR8) has formed side chain intra molecular interactions with the residue Ser422 of flanking region of the hexa peptide at C terminal side (Supplementary figure: 5d). These interactions may hinder the orientation of the mutated (Gly420Glue) IDR for the optimal binding with the polyribosylation enzyme, tankyrase2 and hence binding may not occur. Absence of binding poly ribosylation does not occur and consequently ubiquitination cannot take place to the mutated 3BP2 by RNF16 as it recognizes the ribosylation substrates. The stabilized 3BP2 subsequently hyper activates the SRC kinases and leads to the differentiation of osteoclast, a characteristic of the cellular dysfunction underlying a disease called cherubism (Santanna & Fukai, 2001).

In summary, our studies indicate that deleterious effect of the DCMMs in IDRs is in reducing the conformational heterogeneity that is intrinsic to IDRs thereby restricting their functional plurality. A reduction in the conformational flexibility may mean that the preclusion of certain functionally important conformations for IDRs. Our studies further indicate that the DCMMs stabilize certain conformational states of the IDRs that are not conducive for interactions with their cognate interacting partners thereby hindering their molecular functions.

## Conclusions

The functional impact of the DCMMs on the IDRs was investigated by the MD simulations. From the examples that we considered in this study it is clear that the DCMMs impair the intrinsic conformational heterogeneity of IDRs and furthermore, also appear to preclude the functionally important conformational states at the cost of stabilizing some other conformational states. Although this conclusion has been reached using a small set of IDRs harboring known DCMMs, it is reasonable to believe that this is how the DCMMs affect the IDRs and their functions.

## Supporting information

https://drive.google.com/file/d/1DDYuMBFtfWUvpHI3Atwv1pa0Iu3iResC/view?usp=sharing

https://drive.google.com/file/d/1bDKm_U_2cEfmYBl_A0vwhw1hoWo9u7Ji/view?usp=sharing

## Acknowledgements

HAN and SS together gratefully acknowledge the core grant support of Centre for DNA Fingerprinting and Diagnostic (CDFD) and University of Hyderabad. SS was a recipient of University Grants Commission (UGC) Junior as well as Senior Research Fellowships. He also gratefully acknowledges a financial support from CDFD.

## Declaration of interest statement

Authors declared there is no conflict of interest

## Notes

### Competing Interest Statement

The authors have declared no competing interest.

## References

Borriello, A., Cucciolla, V., Oliva, A., Zappia, V., & Della Ragione, F. (2007). p27Kip1 metabolism: A fascinating labyrinth. Cell Cycle, 6(9), 1053–1061. https://doi.org/10.4161/cc.6.9.4142

Brown, C. J., Johnson, A. K., Dunker, A. K., & Daughdrill, G. W. Evolution and disorder, 21 Current Opinion in Structural Biology § (2011). Elsevier Ltd. https://doi.org/10.1016/j.sbi.2011.02.005

Campen, A., Williams, R. M., Brown, C. J., Meng, J., Uversky, V. N., & Dunker, A. K. (2008). TOP-IDP-scale: a new amino acid scale measuring propensity for intrinsic disorder. Protein and Peptide Letters, 15(9), 956–963. https://doi.org/10.2174/092986608785849164

Cory, G. O. C., Cramer, R., Blanchoin, L., & Ridley, A. J. (2003). Phosphorylation of the WASP-VCA Domain Increases Its Affinity for the Arp2 / 3 Complex and Enhances Actin Polymerization by WASP, 11, 1229–1239.

Davey, N. E., Van Roey, K., Weatheritt, R. J., Toedt, G., Uyar, B., Altenberg, B., … Gibson, T. J. (2012). Attributes of short linear motifs. Molecular BioSystems, 8(1), 268–281. https://doi.org/10.1039/c1mb05231d

Dinauer, M. C., Pierceo, E. A., Erickson, R. W., Muhlebach, T. J., Messnerii, H., Orkint, S. H., … Curnutte, J. T. (1991). Point mutation in the cytoplasmic domain of the neutrophil p22-phox cytochrome b subunit is associated with a nonfunctional NADPH oxidase and chronic granulomatous disease, 88(December), 11231–11235.

Dogan, J., Mu, X., Engström, Å., & Jemth, P. The transition state structure for coupled binding and folding of disordered protein domains, 3 Scientific Reports § (2013). https://doi.org/10.1038/srep02076

Dosztanyi, Z., Chen, J., & Dunker, A. (2006). Disorder and Sequence Repeats in Hub Proteins and Their Implication for Network Evolution. Journal of Proteome …, 5(11), 2985–2995. Retrieved from http://pubs.acs.org/doi/abs/10.1021/pr060171o

Dosztányi, Z., Csizmok, V., Tompa, P., & Simon, I. (2005). IUPred: Web server for the prediction of intrinsically unstructured regions of proteins based on estimated energy content. Bioinformatics, 21(16), 3433–3434. https://doi.org/10.1093/bioinformatics/bti541

Dunker, A. K., Brown, C. J., Lawson, J. D., Iakoucheva, L. M., & Obradovic, Z. Intrinsic disorder and protein function, 41 Biochemistry (John Wiley & Sons) § (2002). https://doi.org/10.1021/bi012159+

Dunker, A. K., Obradovic, Z., Romero, P., Garner, E. C., & Brown, C. J. Intrinsic protein disorder in complete genomes., 11 Genome informatics. Workshop on Genome Informatics § (2000). https://doi.org/10.11234/gi1990.11.161

Dye-terminator, T., Abdul-manan, N., Aghazadeh, B., Liu, G. A., Majumdar, A., & Ouerfelli, O. (1999). Structure of Cdc42 in complex with the GTPase-binding domain of the ‘Wiskott – Aldrich syndrome’ protein, 399(May), 379–383.

Espinoza-Fonseca, L. M. (2009). Leucine-rich hydrophobic clusters promote folding of the N-terminus of the intrinsically disordered transactivation domain of p53. FEBS Letters, 583(3), 556–560. https://doi.org/10.1016/j.febslet.2008.12.060

Fisher, C. K., & Stultz, C. M. Constructing ensembles for intrinsically disordered proteins, 21 Current Opinion in Structural Biology § (2011). Elsevier Ltd. https://doi.org/10.1016/j.sbi.2011.04.001

Guettler, S., Larose, J., Petsalaki, E., Gish, G., Scotter, A., Pawson, T., … Sicheri, F. (2011). Structural basis and sequence rules for substrate recognition by tankyrase explain the basis for cherubism disease. Cell, 147(6), 1340–1354. https://doi.org/10.1016/j.cell.2011.10.046

Haynes, C., Oldfield, C. J., Ji, F., Klitgord, N., Cusick, M. E., Radivojac, P., … Iakoucheva, L. M. Intrinsic disorder is a common feature of hub proteins from four eukaryotic interactomes, 2 PLoS Computational Biology § (2006). https://doi.org/10.1371/journal.pcbi.0020100

Hilarius, P. M., Weening, R. S., Kaulfersch, W., Seger, R. A., Roos, I. I. D., & Verhoeven, A. J. (1994). ‘S6Pro --* Gin Substitution in the Light Chain of Cytochrome bsss of the Human N A D P H Oxidase (p22-phox) Leads to Defective Translocation of the Cytosofic Proteins, 180(December).

Huang, J., Rauscher, S., Nawrocki, G., Ran, T., Feig, M., de Groot, B. L., … MacKerell, A. D. (2017). CHARMM36: An Improved Force Field for Folded and Intrinsically Disordered Proteins. Biophysical Journal, 112(3), 175a–176a. https://doi.org/10.1016/j.bpj.2016.11.971

Iakoucheva, L. M., Brown, C. J., Lawson, J. D., Obradović, Z., & Dunker, A. K. Intrinsic disorder in cell-signaling and cancer-associated proteins, 323 Journal of Molecular Biology § (2002). https://doi.org/10.1016/S0022-2836(02)00969-5

Iakoucheva, L. M., Radivojac, P., Brown, C. J., O’Connor, T. R., Sikes, J. G., Obradovic, Z., & Dunker, A. K. (2004). The importance of intrinsic disorder for protein phosphorylation. Nucleic Acids Research, 32(3), 1037–1049. https://doi.org/10.1093/nar/gkh253

Jones, D. T., & Cozzetto, D. DISOPRED3: Precise disordered region predictions with annotated protein-binding activity, 31 Bioinformatics § (2015). https://doi.org/10.1093/bioinformatics/btu744

Kar, S., Sakaguchi, K., Shimohigashi, Y., Samaddar, S., Banerjee, R., Basu, G., … Roy, S. (2002). Effect of phosphorylation on the structure and fold of transactivation domain of p53. Journal of Biological Chemistry, 277(18), 15579–15585. https://doi.org/10.1074/jbc.M106915200

Levaot, N., Voytyuk, O., Dimitriou, I., Sircoulomb, F., Chandrakumar, A., Deckert, M., … Rottapel, R. (2011). Loss of Tankyrase-mediated destruction of 3BP2 is the underlying pathogenic mechanism of cherubism. Cell, 147(6), 1324–1339. https://doi.org/10.1016/j.cell.2011.10.045

Lowry, D. F., Stancik, A., Shrestha, R. M., & Daughdrill, G. W. (2008). Modeling the accessible conformations of the intrinsically unstructured transactivation domain of p53. Proteins: Structure, Function and Genetics, 71(2), 587–598. https://doi.org/10.1002/prot.21721

Luthi, J. N., Gandhi, M. J., & Drachman, J. G. (2003). X-linked thrombocytopenia caused by a mutation in the Wiskott-Aldrich syndrome (WAS) gene that disrupts interaction with the WAS protein (WASP)-interacting protein (WIP), 31, 150–158.

Michel Espinoza-Fonseca, L., Ilizaliturri-Flores, I., & Correa-Basurto, J. (2012). Backbone conformational preferences of an intrinsically disordered protein in solution. Molecular BioSystems, 8(6), 1798. https://doi.org/10.1039/c2mb00004k

Obradovic, Z., Uversky, V. N., Manuscript, A., Terms, T., Regions, D., Uversky, V. N., & Obradovic, Z. NIH Public Access, 6 Proteome § (2008). https://doi.org/10.1021/pr060394e.Functional

Ogura, K., Nobuhisa, I., Yuzawa, S., Takeya, R., Torikai, S., Saikawa, K., … Inagaki, F. (2006). NMR Solution Structure of the Tandem Src Homology 3 Domains of p47 phox Complexed with a p22 phox -derived, 281(6), 3660–3668. https://doi.org/10.1074/jbc.M505193200

Oldfield, C. J., Meng, J., Yang, J. Y., Yang, M. Q., Uversky, V. N., & Dunker, A. K. Flexible nets: disorder and induced fit in the associations of p53 and 14-3-3 with their partners, 9 BMC Genomics § (2008). https://doi.org/10.1186/1471-2164-9-S1-S1

Patil, A., & Nakamura, H. (2006). Disordered domains and high surface charge confer hubs with the ability to interact with multiple proteins in interaction networks. FEBS Letters, 580(8), 2041–2045. https://doi.org/10.1016/j.febslet.2006.03.003

Qin, B. Y., Liu, C., Srinath, H., Lam, S. S., Correia, J. J., Derynck, R., & Lin, K. (2005). Crystal structure of IRF-3 in complex with CBP. Structure, 13(9), 1269–1277. https://doi.org/10.1016/j.str.2005.06.011

Radivojac, P., Vacic, V., Haynes, C., Cocklin, R. R., Mohan, A., Heyen, J. W., … Iakoucheva, L. M. (2011). Identification, Analysis and Prediction of Protein Ubiquitination Sites. Proteins, 78(2), 365–380. https://doi.org/10.1002/prot.22555.Identification

Santanna, C., & Fukai, N. (2001). Ng0601_125, 28(june).

Schaefer, C., Schlessinger, A., & Rost, B. (2010). Protein secondary structure appears to be robust under in silico evolution while protein disorder appears not to be. Bioinformatics, 26(5), 625–631. https://doi.org/10.1093/bioinformatics/btq012

Singh, G. P., Ganapathi, M., Sandhu, K. S., & Dash, D. (2006). Intrinsic unstructuredness and abundance of PEST motifs in eukaryotic proteomes. Proteins: Structure, Function and Genetics, 62(2), 309–315. https://doi.org/10.1002/prot.20746

Sugase, K., Dyson, H. J., & Wright, P. E. Mechanism of coupled folding and binding of an intrinsically disordered protein, 447 Nature § (2007). https://doi.org/10.1038/nature05858

Turjanski, A. G., Gutkind, J. S., Best, R. B., & Hummer, G. (2008). Binding-induced folding of a natively unstructured transcription factor. PLoS Computational Biology, 4(4), 1–11. https://doi.org/10.1371/journal.pcbi.1000060

Uversky, V. N., & Dunker, A. K. Understanding protein non-folding, 1804 Biochimica et Biophysica Acta - Proteins and Proteomics § (2010). Elsevier B.V. https://doi.org/10.1016/j.bbapap.2010.01.017

Uversky, V. N., Gillespie, J. R., & Fink, A. L. (2000). Why are “natively unfolded” proteins unstructured under physiologic conditions? Proteins: Structure, Function and Genetics, 41(3), 415–427. https://doi.org/10.1002/1097-0134(20001115)41:3<415::AID-PROT130>3.0.CO;2-7

Uversky, V. N., Oldfield, C. J., & Dunker, A. K. (2005). Showing your ID: Intrinsic disorder as an ID for recognition, regulation and cell signaling. Journal of Molecular Recognition, 18(5), 343–384. https://doi.org/10.1002/jmr.747

Uversky, V. N., Oldfield, C. J., & Dunker, A. K. Intrinsically Disordered Proteins in Human Diseases: Introducing the D 2 Concept, 37 Annual Review of Biophysics § (2008). https://doi.org/10.1146/annurev.biophys.37.032807.125924

Vacic, V., & Iakoucheva, L. M. (2012). Disease mutations in disordered regions—exception to the rule? Mol. BioSyst., 8(1), 27–32. https://doi.org/10.1039/C1MB05251A

Vucetic, S., Brown, C. J., Dunker, A. K., & Obradovic, Z. Flavors of protein disorder, 52 Proteins: Structure, Function and Genetics § (2003). https://doi.org/10.1002/prot.10437

Ward, J. J., Sodhi, J. S., McGuffin, L. J., Buxton, B. F., & Jones, D. T. Prediction and Functional Analysis of Native Disorder in Proteins from the Three Kingdoms of Life, 337 Journal of Molecular Biology § (2004). https://doi.org/10.1016/j.jmb.2004.02.002

Ward, Jonathan J., McGuffin, L. J., Bryson, K., Buxton, B. F., & Jones, D. T. The DISOPRED server for the prediction of protein disorder, 20 Bioinformatics § (2004). https://doi.org/10.1093/bioinformatics/bth195

Wright, P. E., & Dyson, H. J. Intrinsically unstructured proteins: re-assessing the protein structure-function paradigm, 293 Journal of Molecular Biology § (1999). https://doi.org/10.1006/jmbi.1999.3110

Xue, B., Dunbrack, R. L., Williams, R. W., Dunker, A. K., & Uversky, V. N. (2010). PONDR-FIT: A meta-predictor of intrinsically disordered amino acids. Biochimica et Biophysica Acta - Proteins and Proteomics (Vol. 1804). https://doi.org/10.1016/j.bbapap.2010.01.011

