## Supplementary material for "Effect of Disease Causing Missense Mutations on Intrinsically Disordered Regions in Proteins": https://drive.google.com/file/d/1DDYuMBFtfWUvpHI3Atwv1pa0Iu3iResC/view?usp=sharing

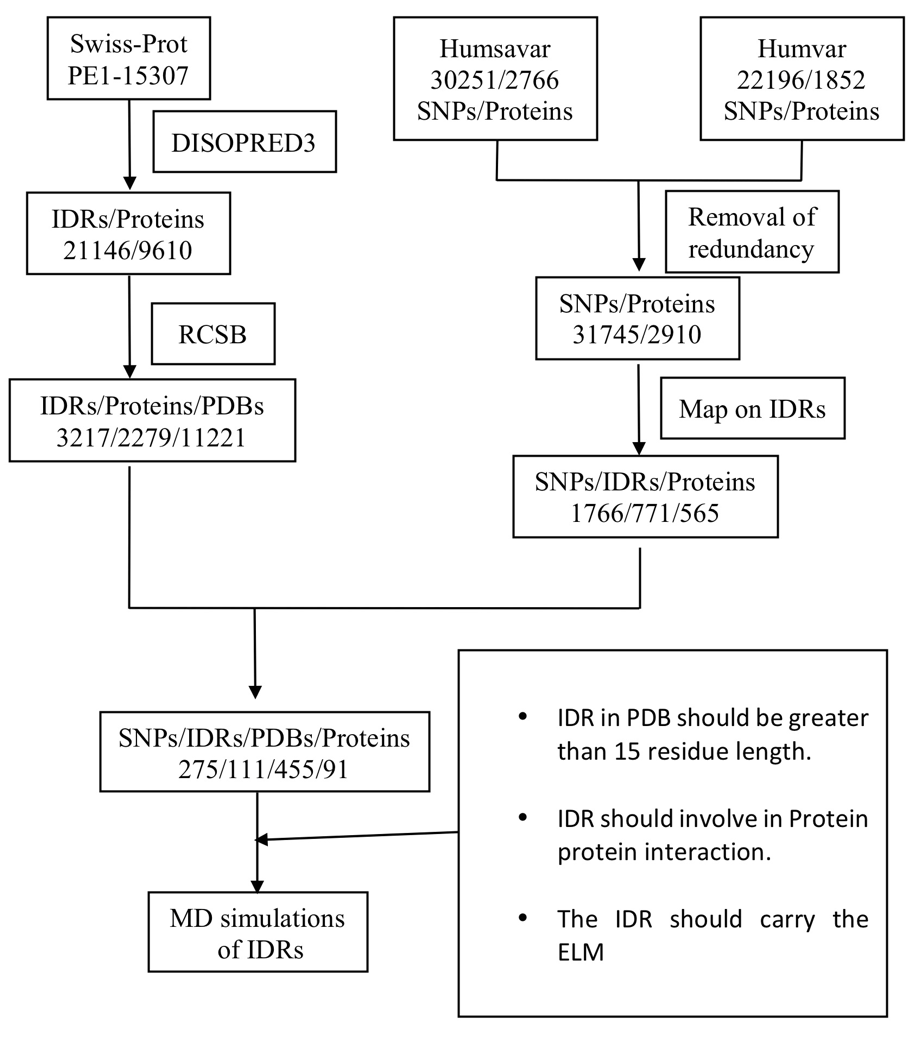

**Supplementary figure 1:** We considered proteins which have evidence of the existence at protein level (PE1) from Swiss-Prot and supplied them to disopred3 to predict their intrinsic disorderedness. We considered long intrinsically disordered regions (IDRs) and retrieved the structural information for the same IDRs from the protein data bank, RCSB and considered only those having X-ray and NMR structures. On the other hand, we downloaded disease causing missense mutations from humsavar and humvar and removed the duplicates and mapped these disease SNPs onto the IDRs. Further, we retrieved the structures for IDRs with disease SNPs from protein data bank (PDB, RCSB). Based on the criteria (length of IDR, involve in PPI and experimentally validated ELM) we selected three IDRs for the case study of MD simulations.

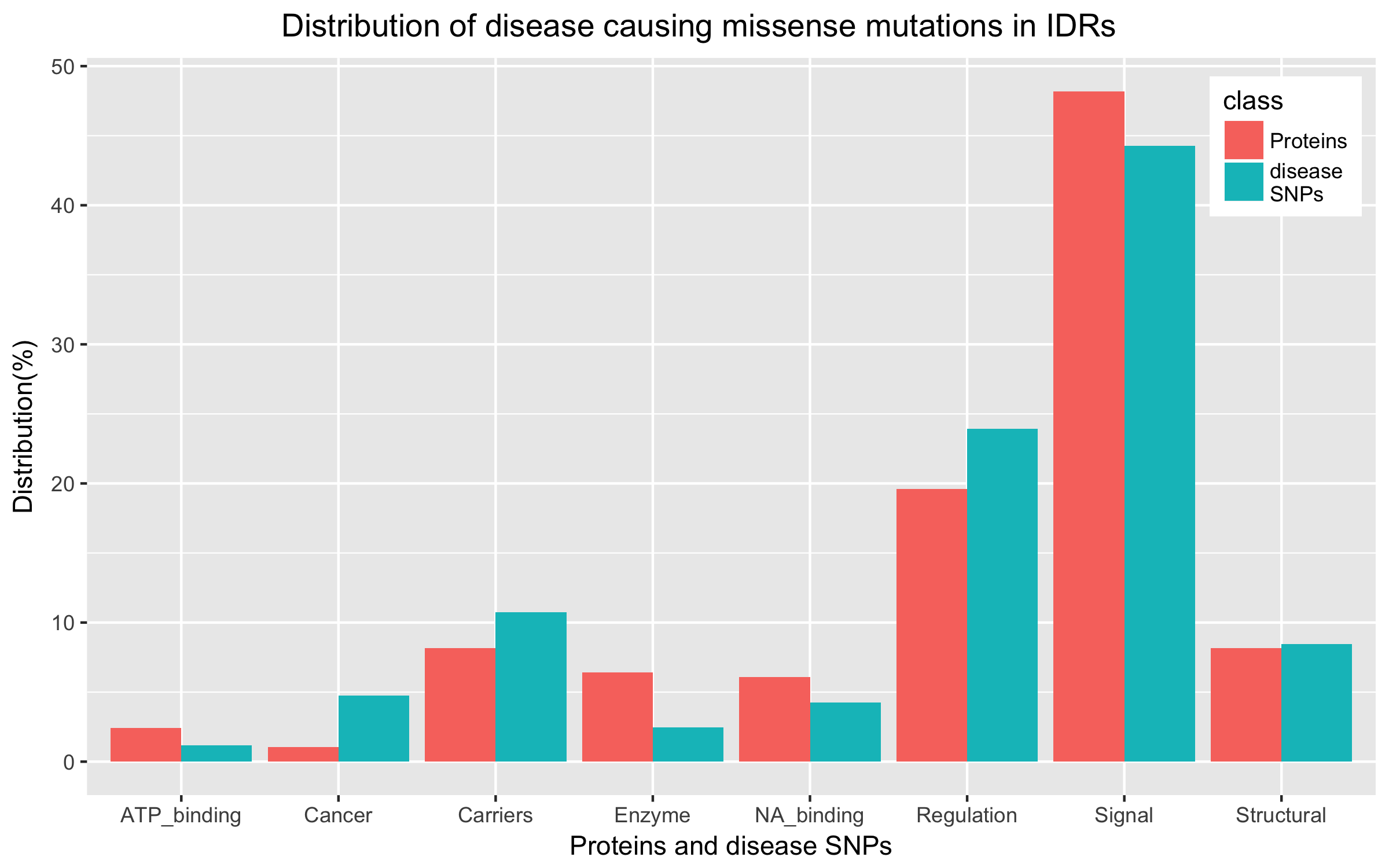

**Supplementary figure 2:** Most of the disease causing missense mutations (DCMMs) harboring IDRs are parts of the proteins involved in signaling pathways. The other proteins harboring the DCMMs are regulation (transcription and translation). kinases, GTPases, scaffolding proteins and other downstream effector proteins. The mutations in the regulation may affect DNA binding properties. The structural proteins include cytoskeletal elements and extracellular proteins. The IDRs in the structural proteins might mediate the protein-protein interactions to give strength. The DCMMs in these IDRs could hamper their interactions. The topological domains of transporters and channels are predicted to be intrinsically disordered and some of the disease mutations mapped in these regions and could affect the binding of the transported molecules. Nucleic acid-binding proteins, some of the enzymes and cell–adhesion proteins harbor DCMMs in their IDRs.

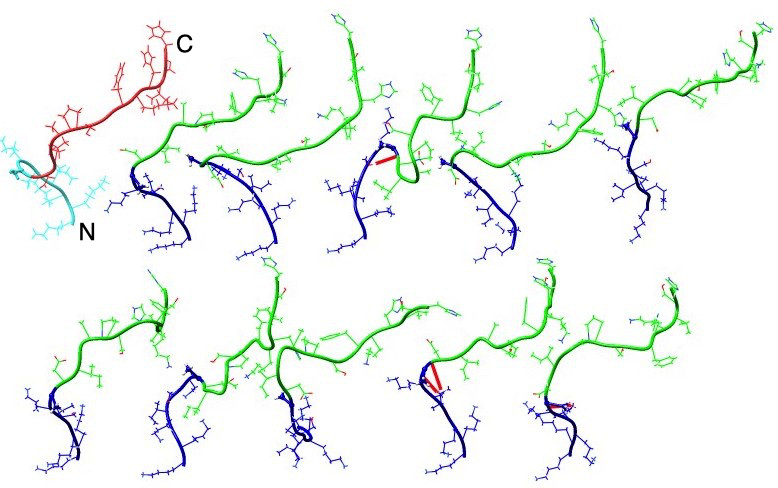
 **Supplementary figure 3a:** The WT conformational states of CRIB IDR are comprised of transient turns, bends and loops. They form a few main chain interactions (shown in red). The residues are shown in the ball and stick model. The blue colored region represents the N terminal region of the CRIB IDR and this region harbors the mutation site. The red and cyan colored structure in the image is the IDR of the NMR structure. The letters N and C respectively represent N-terminal and C-terminal ends of the IDRs. All the conformations are in the same orientation. N and C represent the N terminal and C terminal side of IDR respectively.

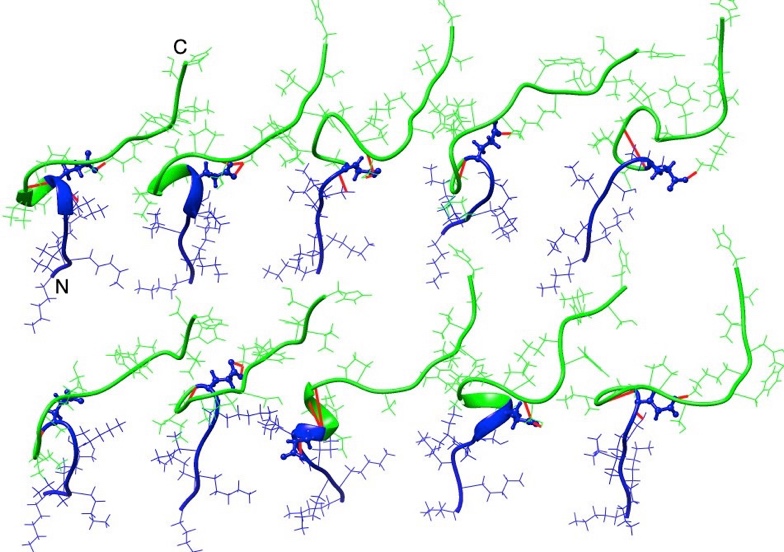

**Supplementary figure 3b:** The Glu residue in A236E mutant structure in the CRIB IDR is involved in hydrogen bond interactions and stabilizes the helix and turns. The MT residue is shown in the ball and stick model and the red-coloured lines represent interactions. The letters N and C respectively represent N-terminal and C-terminal ends of the IDRs. All the conformations are in the same orientation. N and C represent the N terminal and C terminal side of IDR respectively.

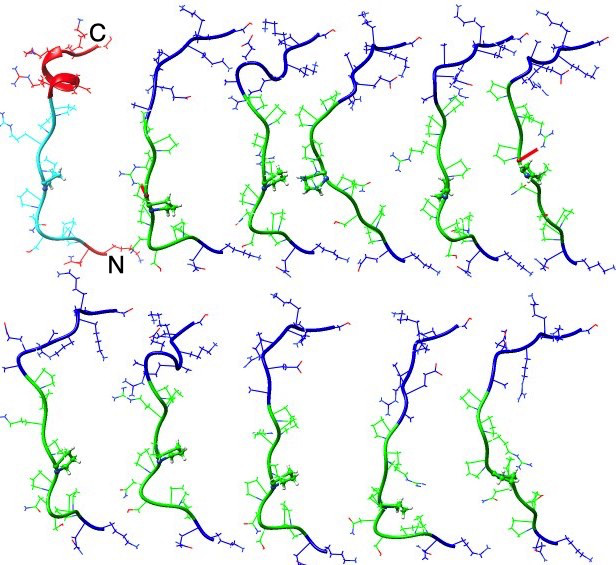

**Supplementary figure 4a:** The WT PRR IDR does not form many interactions. The green colored region represents the binding region with SH3 domain of p47 and it adopts polyproline type II helix. Most of the distinct conformational substates of WT adopt the polyproline type II helix during the simulations. The red and cyan colored structure in the image is the IDR of the NMR structure. N and C represent the N terminal and C terminal side of IDR respectively.

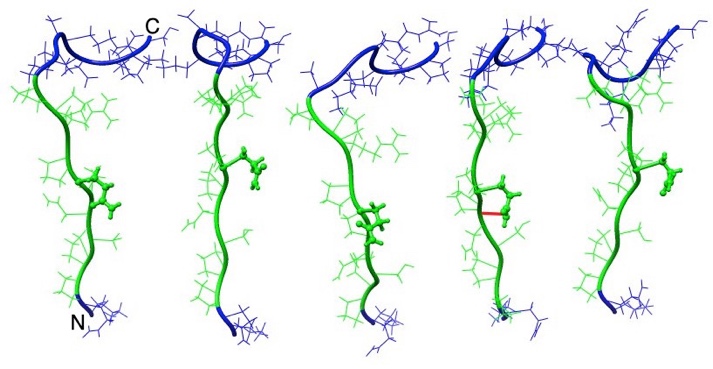

**Supplementary figure 4b:** The MT PRR IDR also does not form intra molecular interaction in the distinct conformational substates except one (IDR4) during the simulations. Red coloured line represents the intra molecular interaction. However, the N-terminal region adopts a folded structure as compared with the WT. N and C represent the N terminal and C terminal side of IDR respectively.

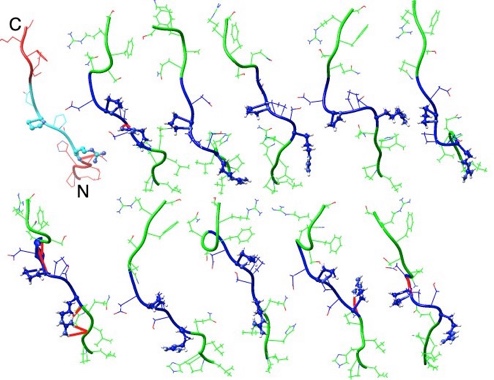
 **Supplementary figure 5a:** Distinct conformational substates of the WT of TRM IDR. The blue region represents the hexa peptide which is crucial for the binding with the tankyrase2 in extended conformation. The ball and stick residues represent the WT residues such as R415, P418, and G420. R415 formed few side chain interactions (red coloured lines) in distinct conformational substates (IDR2, IDR7 and IDR 10). The red and cyan colored structure in the image represents the IDR of the crystal structure. N and C represent the N terminal and C terminal side of IDR respectively.

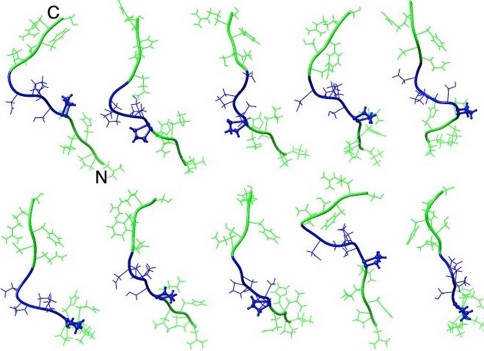

**Supplementary figure 5b:** The MT residue R415P (ball and stick model) in the TRM IDR introduced the kink in the binding regions (blue colored) in all the distinct conformational substates and thereby disrupted the extended conformation of the binding peptide. N and C represent the N terminal and C terminal side of IDR respectively.

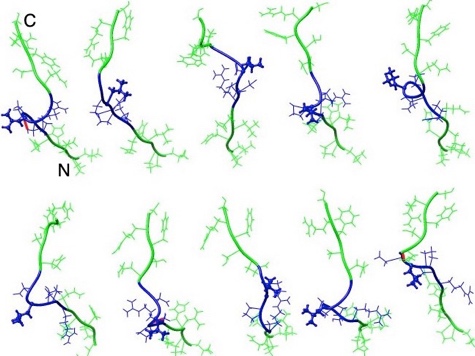
**Supplementary figure 5c:** The P418L (ball and stick model) mutation in the TRM IDR formed main chain interactions in the distinct conformational substates (IDR1, IDR5, and IDR8) and stabilized them during the simulations. These three clusters encompassed the one-third of the simulation time (30.5ns). Because of mutation, the binding region (blue color) of TRM IDR adopts turns and bends which disrupts the extended conformation of the binding region (blue color). N and C represent the N terminal and C terminal side of IDR respectively.

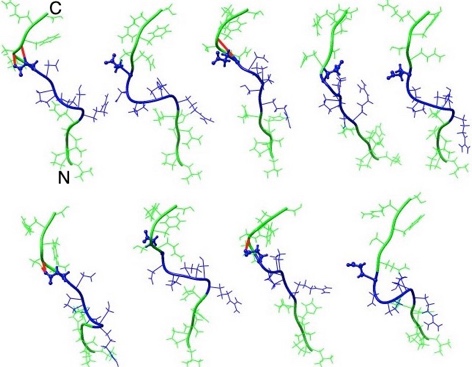
**Supplementary figure 5d:** The mutation G420E (ball and stick model) established the side chain interactions (red coloured lines) in the distinct conformational substates of 1, 6, and 8. The clusters are represented by distinct conformational substates (IDR1, IDR6, and IDR9) occupied 40% of the time during the simulations. The mutation G420E adopts turns in the binding region (blue color) of the TRM IDR and there disrupted the extended conformation which is required for the interaction. N and C represent the N terminal and C terminal side of IDR respectively.

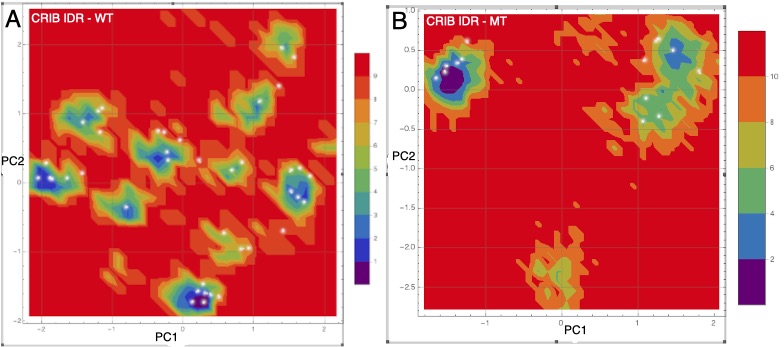

**Supplementary figure 6:** The figure shows the energy landscape of the CRIB IDR. A and B represents the FEL of WT and MT IDR respectively. The energy local minima of FEL are shown as blue colored regions. The figure also shows location of various conformations of WT and MT IDRs in relation to the local energy minima. The conformations are shown as white dots.

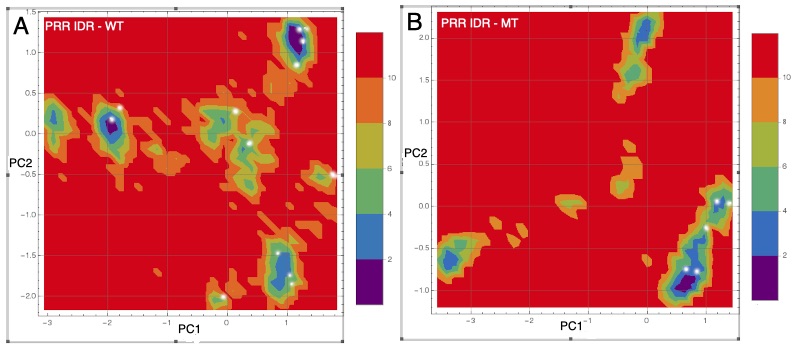

**Supplementary figure 7:** The figure shows the FEL of the PRR IDR. A and B represents the FEL of WT and MT IDR respectively. The energy local minima of FEL are shown as blue colored regions. The figure also shows location of various conformations of WT and MT IDRs in relation to the local energy minima. The conformations are shown as white dots.

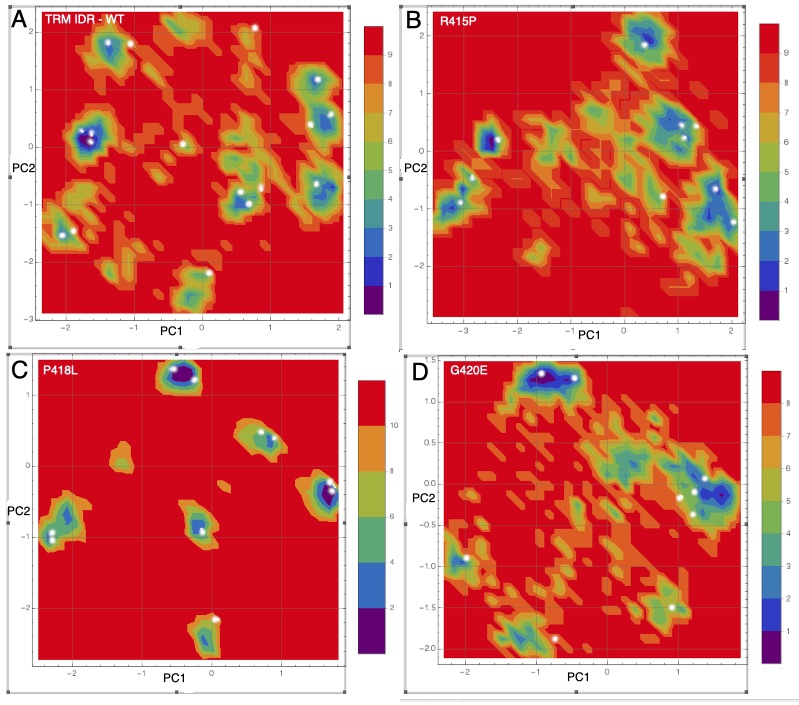

**Supplementary figure 8:** The figure shows the FEL of the TRM IDR. A bin represents the WT and B to D represent the FEL of R415P, P418L and G420E respectively. The energy local minima of FEL are shown as blue colored regions. The figure also shows location of various conformations of WT and MT IDRs in relation to the local energy minima. The conformations are shown as white dots.
