## Supplementary material for "Effect of Disease Causing Missense Mutations on Intrinsically Disordered Regions in Proteins": https://drive.google.com/file/d/1bDKm_U_2cEfmYBl_A0vwhw1hoWo9u7Ji/view?usp=sharing

**Supplementary table 1:** Essential information of the IDRs selected for MD simulations

| **IDR ID (PDB)** | | **No. of atoms in IDR** | **No.of solvent molecules** | **Counter ions** | **EM steps** |
| --- | --- | --- | --- | --- | --- |
| **1CEE-B :18** | WT | 314 | 3759 | 4 cl^-^ | 359 |
|  | A236E | 319 | 3715 | 3 cl^-^ | 594 |
| **1WLP-A : 20** | WT | 317 | 2939 | 4 cl^-^ | 277 |
|  | P156Q | 320 | 3019 | 4 cl^-^ | 239 |
| **3TWR-H : 16** | WT | 254 | 3509 | 1 cl^-^ | 225 |
|  | R415P | 244 | 3431 | 0 | 210 |
|  | P418L | 259 | 3484 | 1 cl^-^ | 135 |
|  | G420E | 262 | 3490 | 0 | 130 |

**Supplementary table 2:** The RMSD values of the hexa peptide of the distinct conformational substates of WT and MT IDRs with respect to the same hexa peptide of the crystal structure of the TRM IDR present in the 3BP2. Particularly the MT IDRs such as R415P and P418L highly deviated from the hexa peptide of the crystal structure during the simulations.

| **Distinct IDRs** | **RMSD (nm) of hexa peptide** | | | |
| --- | --- | --- | --- | --- |
|  | **WT** | **R415P** | **P418L** | **G420E** |
| 1 | 0.06 | 0.36 | 0.36 | 0.05 |
| 2 | 0.11 | 0.31 | 0.19 | 0.17 |
| 3 | 0.08 | 0.25 | 0.38 | 0.18 |
| 4 | 0.15 | 0.26 | 0.34 | 0.07 |
| 5 | 0.10 | 0.28 | 0.37 | 0.16 |
| 6 | 0.22 | 0.20 | 0.32 | 0.05 |
| 7 | 0.06 | 0.34 | 0.32 | 0.17 |
| 8 | 0.10 | 0.30 | 0.19 | 0.08 |
| 9 | 0.03 | 0.29 | 0.35 | 0.13 |
| 10 | 0.25 | 0.28 | 0.15 | - |
| 11 | 0.17 | - | - | - |
| 12 | 0.07 | - | - | - |
| 13 | 0.08 | - | - | - |
| 14 | 0.21 | - | - | - |
| 15 | 0.07 | - | - | - |
| 16 | 0.08 | - | - | - |
| 17 | 0.12 | - | - | - |
